# Adaptation to high pressure; insights from the genome of an evolved *Escherichia coli* strain with increased piezotolerance

**DOI:** 10.1101/2024.09.24.613341

**Authors:** Marco N. Allemann, Stacey Ma, Terra Sztain, Thomas G. Bartholow, Ian P.G. Marshall, J. Andrew McCammon, Michael Burkart, Eric E. Allen, Douglas H. Bartlett, Angeliki Marietou

## Abstract

Pressure is a key environmental parameter that influences the activity and distribution of microbial life on our planet. Despite its key role there is still not a definitive list of essential genes for microbial adaptation to life under increasing pressure. In this study we used a previously characterized *Escherichia coli* strain (AN62) evolved to grow at pressure (60 MPa) non-permissive to the parental strain and performed comparative genomics in order to identify the genome-level adaptations that might allowed the observed pressure-adapted phenotype. We identified 18 mutations in total of which 3 mutations were present in both the parental and evolved strain, 3 mutations were only present in the parental strain, and 12 mutations were observed only in the evolved AN62 strain. Among the characterized mutations we observed a point mutation in the acyl carrier protein (*acp*P^V43G^). Complementation experiments revealed that the observed V43G mutation in AcpP is responsible for increased levels of cis-vaccenic acid but is not alone responsible for the pressure adapted phenotype. Further molecular dynamics and docking simulations suggested that the V43G mutation promoted stronger binding of the AcpP protein to partner enzymes of the fatty acid biosynthesis pathway involved in fatty acid unsaturation.

**Data Summary:** *Escherichia* coli reads from the parental and evolved strain have been deposited in the Sequence Read Archive (SRA) under accession number RJNA600359. All software used in the bioinformatic analysis is publicly available.

**Impact Statement:** Pressure is a key environmental parameter. Two-thirds of our planet is covered by oceans with an average depth of 3800m, which means that the majority of the marine life experiences deep sea conditions. Our results offer a list of gene mutations that could contribute to an improved pressure growth phenotype in *Escherichia coli*, offering a unique insight on the genome level adaptations that might contribute to high pressure adaptation.

## Introduction

Pressure is a key environmental parameter that influences life in the deep sea. As we travel from the sea surface down the water column pressure increases by 1MPa for every 100 m. Organisms with optimal growth rates above 0.1 MPa (atmospheric pressure) are defined as piezophiles (from the Greek “piezo”, meaning “to press”) and span a wide range of bacterial and archaeal lineages. Over 70% of the Earth’s biosphere is under high pressure (Picard and Daniel, 2013). The pleotropic effects of pressure have been described extensively in the model mesophilic bacterium *Escherichia coli*. Increasing pressure progressively “shuts downs” key cellular processes starting with motility at 10 MPa and substrate transport at 26 MPa (Meganathan and Marquis, 1973; Landau, 1967). Pressures between 20 and 50 MPa affect cell division resulting in elongation of the cells and the characteristic piezosensitive filamentous morphology (Zobell and Cobet, 1964). DNA synthesis is in turn abolished at 50 MPa, followed by translation at 60 MPa and transcription at 77 MPa (Yayanos and Pollard, 1969). Our understanding of the mechanisms adopted by piezophiles in order to adapt and thrive at increasing pressures is limited but the handful of studies available suggest that it involves adjusting gene and protein expression and the production of osmolytes (Simonato et al., 2006; Campanaro et al., 2012; Michoud and Jebbar, 2016). The most well-characterized response of piezophiles to pressure is their ability to increase the proportion of unsaturated fatty acids in their lipids to preserve membrane fluidity (Allen et al., 1999).

Biosynthesis of fatty acids in *Escherichia coli* occurs via the well-studied Type II fatty acid synthase (FAS) pathway (Cronan and Rock, 2008; Magnuson et al., 1993). The acyl-carrier protein (AcpP) is a key protein in fatty acid synthesis as it transports the growing fatty acid chain between the different enzymes involved in fatty acid biosynthesis (Nguyen et al., 2014). During fatty acid biosynthsis the acyl group is covalently attached to the post-translationally modified AcpP through a thioester linkage (Cronan and Rock, 2008). Given this critical role *acp*P is considered an essential gene and as one of the most abundant proteins in *E.coli*. AcpP has also been shown to interact with a variety of other proteins not related to fatty acid biosynthesis (Cronan and Rock, 2008). One well-characterized example is the interaction with SpoT, a bifunctional ppGpp synthetase/hydrolase that mediates the stringent response (Angelini et al., 2012; Battesti and Bouveret, 2006; Seyfzadeh, 1993). It has been suggested that SpoT monitors the metabolic state of the cell through this interaction and adjusts the ppGpp pool in response to alterations of the acyl-AcpP pool (Battesti and Bouveret, 2006). During the fatty acid biosynthetic process, AcpP interacts with at least 12 different enzymes of the FAS pathway including FabA and FabF (Nguyen et al., 2014). FabF is a ketoacyl synthase that is responsible for the elongation of C14:1 and catalyses the last step to vaccenic acid (C18:1) (Garwin and Cronan, 1980). FabA acts as a dehydratase and is essential for the synthesis of unsaturated fatty acids (Beld et al., 2015).

Piezophiles have been isolated and characterized within the Thermococci, Methanococci, Methanopyri, Bacilli, Actinobacteria, Alphaproteobacteria, Deltaproteobacteria, and Thermotogae with the majority of the isolates belonging to the Gammaproteobacteria class (Eloe et al., 2011). The molecular blueprint that allows these piezophilic isolates to thrive under high pressure conditions remains yet unknown. Comparative genomics and other “omics” approaches of the very few sequenced obligate piezophilic strains highlighted several pressure-responsive genes involved in translation, chemotaxis, motility, and energy metabolism (Lauro et al., 2014; Michoud and Jebbar, 2016; Zeng et al., 2019). Evidence from the presence of polyphyletic clusters of piezophilic strains is indicative of various separate evolutionary events towards high pressure adaptation over time (Peoples and Bartlett, 2017). Comparative genomics of shallow and deep sea *Photobacterium* spp. suggest that horizontal gene transfer might play a key role in the evolution of deep-sea adapted strains (Lauro et al., 2014).

Following 500 generations of adaptive laboratory evolution experiments (ALE) we have previously isolated an *E.coli* strain with the ability to grow at pressure non-permissive to the parental strain (Marietou et al., 2015). The strain was able to grow at 60 MPa and change the proportion of unsaturated fatty acids in response to high pressure, a fundamental property of most piezotolerant and piezophilic strains. The aim of this analysis was to identify the underlying mutations that likely contributed to the improved pressure adapted growth phenotype of the evolved strain using whole genome sequencing and comparative genomics. Furthermore, a combination of genetic experiments and simulations were employed in order to investigate the role towards pressure adaptation of an *acp*P mutation identified in the evolved strain.

## Methods

### Comparative genomics

Genomic DNA was extracted from three biological samples for each strain (parental and evolved strain) using the DNeasy Blood & Tissue Kit (Qiagen) following the “Gram Negative Bacteria” protocol. Libraries were prepared using the Nextera XT kit (Illumina, USA) and sequencing was carried out on the Illumina MiSeq platform with a paired-end 300-bp MiSeq v3 reagent kit. The IMG/MER platform (Chen et al., 2019) was used to search (Find Genes> BLAST) the genomes of known piezophilic strains for proteins that were closely related (above 30% identity) to sequences of interest from the *E.coli* genome. The selected amino acid sequences were exported, aligned using Clustal Omega (Sievers et al., 2011), and edited manually. Mapping and identification of SNPs was carried out using Snippy (https://github.com/tseemann/snippy).

### Culturing and strain construction

*Escherichia coli* strains were grown at 37°C in Luria-Bertani (LB) liquid or solid media with addition of 15g/L of agar. The antibiotics ampicillin (100µg/ml) and chloramphenicol (15µg/ml) were added at the indicated concentrations where appropriate. For high-pressure growth, strains were grown in heat-sealed transfer bulbs in stainless steel pressure vessels as described previously (Marietou et al., 2015). A growth indicator; 2, 3, 5-tetrazolium chloride (TTC) was included at 0.005% (w/vol) for visual representation of growth in the high-pressure bulbs. TTC is colourless but in the presence of microbial respiration it is converted to TPF (1,3,5-triphenylformazan) that turns the culture medium violet red.

Strains with various *acp*P alleles were constructed using the lambda red recombineering method (Datsenko and Wanner, 2000) and inserting a chloramphenicol resistance cassette into the nearby *ycf*H gene in both MG1655 (parental) and AN62 (evolved) strains . Once verified the closely linked *acp*P alleles were transfer between MG1655 and AN62 using P1 phage transduction. Chloramphenicol resistant colonies were screened by PCR for the presence of the expected *acp*P allele using *acp*P specific primers (Table 1). PCR products were sequenced by Sanger sequencing (Eton Bioscience, Inc. San Diego, CA, USA) to confirm the sequence of the specific *acp*P allele. The strains and plasmids used in this study are listed in Table 2.

**Table 1.**
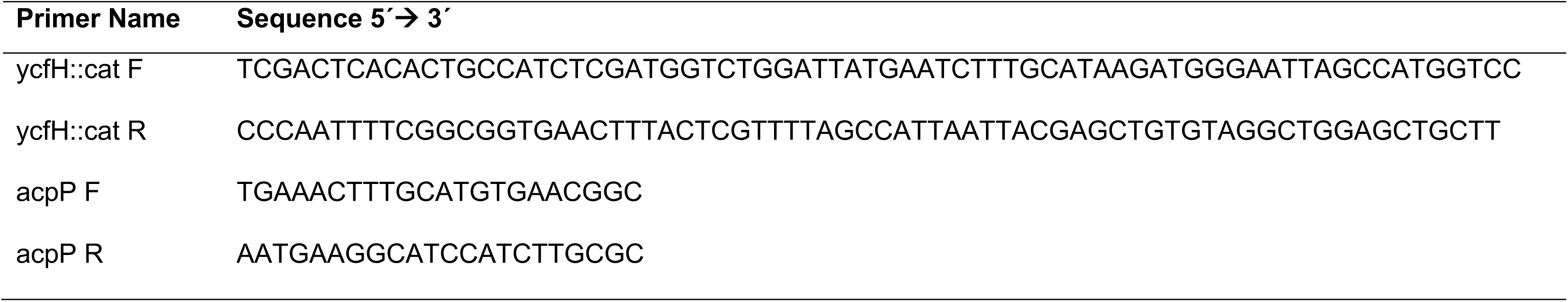
List of primers used in this study.

**Table 2.**
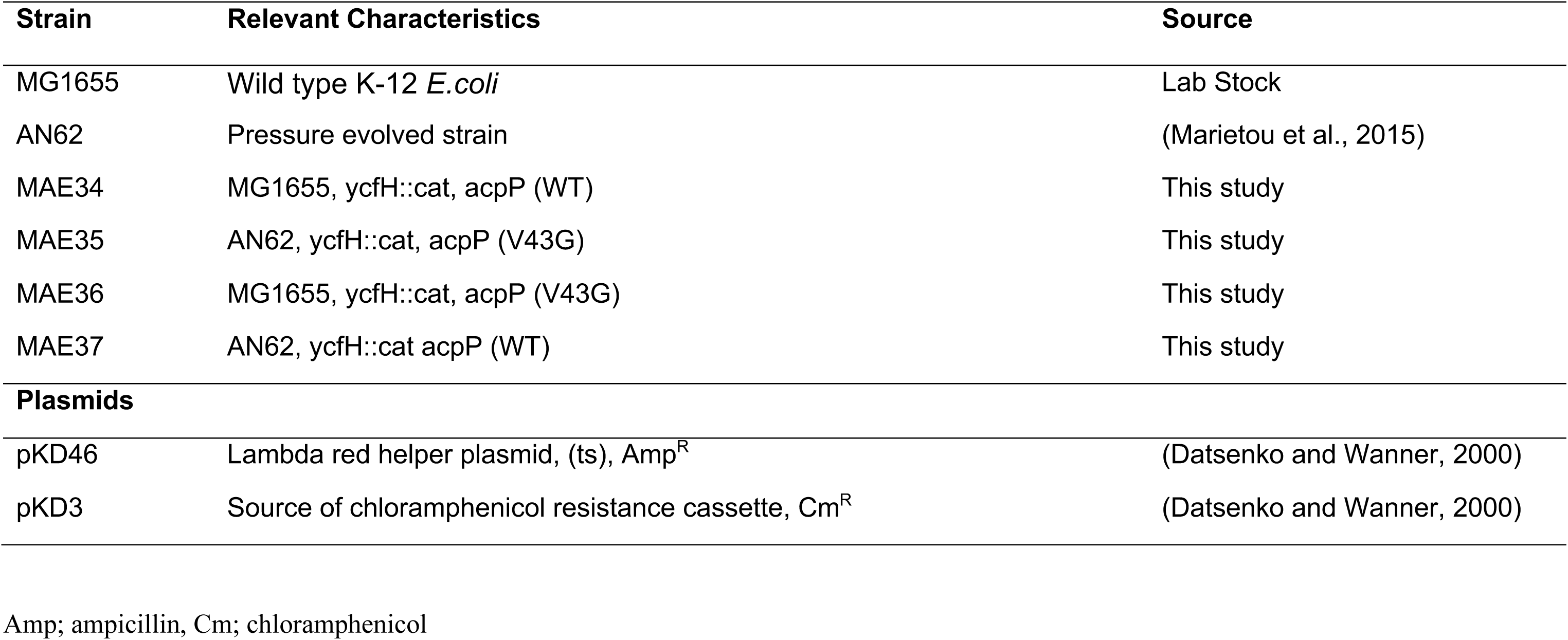
List of *E. coli* strains and plasmids used in this study.

### Fatty acid analysis

For fatty acid analysis, batch cultures (25mL) were grown to late log phase in LB and harvested by centrifugation. Cells pellets were lyophilized and fatty acids methyl esters were prepared and analysed by gas chromatography mass spectrometry (GC-MS) as described previously (Allemann and Allen, 2018).

### Molecular dynamics (MD) simulations

Structures were prepared as described previously (Sztain et al., 2019). Starting from the crystal structure of heptanoyl-AcpP PDB ID 2FAD (Roujeinikova et al., 2007) using chain A, additional carbons and functional groups were added using the Avogadro molecular modeling program (Hanwell et al., 2012) For the 9Z-hexadecenoyl-AcpP, The position of the hexadecanoyl group was obtained from a previous (Sztain et al., unpublished) 500 ns simulation to equilibrate the position of the modeled chain. The cis unsaturation at position 9 was then introduced using the Avogadro molecular modeling program (Hanwell et al., 2012). The AcpP structure contains a conserved serine residue, which is post-translationally modified by a phosphopeantheine arm. The acyl substrates are then covalently attached to the phosphopantetheine via a thioester bond. Therefore, forcefield parameters were generated to treat the ligand as a non-standard amino acid residue. The acyl chain, phosphopantetheine, and serine, were capped with N-methyl and C-acetyl groups for calculation of partial atomic charges according to the restrained electrostatic potential (RESP) method (Bayly et al., 1993). This was calculated using the Hartree-Fock 6-31G(d) method in Gaussian 09. Antechamber was used to generate GAFF (Wang et al., 2004) parameters for the non-proteogenic portion, and Amber ff14SB (Maier et al., 2015) for the serine portion.

Each structure was solvated with an isometric TIP3P water box with at least 10 Å between the protein and edges of the box. The simulation cell was neutralized by adding Na^+^ counterions and additional Na^+^ and Cl^-^ atoms to achieve an ionic strength of 150 mM. Standard minimization, heating, and equilibration were carried out as follows. Minimization was carried out in two steps, with an initial solvent minimization for up to 10,000 cycles with protein residues restrained by a force constant of 500 kcal•mol–1•Å–2 followed by total system minimization for up to 400,000 cycles. Heating was carried out slowly, reaching 310 K over 350 ps. Equilibration was carried out for 1 ns. All production simulations were carried out using AMBER 18 pmemd.cuda (Case et al., 2005; Salomon-Ferrer et al., 2013) at 310 K using NPT ensemble with Monte Carlo barostat and temperature control by Langevin thermostat with a collision frequency of 1 ps-1. Simulations were carried out with 2 fs timesteps and periodic boundary conditions. All nonpolar bonds involving hydrogens and TIP3P water molecules constrained using the SHAKE algorithm (Ryckaert et al., 1977). The Particle Mesh Ewald (PME) method was used to treat long-range electrostatic interactions using a 10.0 Å cutoff for nonbonded interactions. Each acyl-AcpP system was subjected to three independent 500 ns MD simulations. Trajectories were analyzed using CPPTRAJ (Roe and Cheatham, 2013) and graphs were generated in Jupyter Notebooks (Kluyver et al., 2016) using matplotlib (Hunter, 2007). Structures were generated using PyMOL 2.1.1 ( The PyMOL Molecular Graphics System, Version 2.0 Schrödinger, LLC). Pocket volume was calculated using POVME 2.0 (Durrant et al., 2014) from 300 structures obtained from hierarchical clustering based on RMSD from initial structure.

### Docking Calculations

All docking studies were performed using the ICM-Pro docking suite (Abagyan and Totrov, 1994; Abagyan et al., 1994). The FabF (2GFW) and FabA (1MKB) partner protein structures were acquired from the Protein Data Bank (http://www.rcsb.org) (Berman et al., 2000). Both partner proteins were docked as dimers in order to properly recreate the interaction in solution. The ICM quickflood procedure (Fernández-Recio et al., 2003) was used to generate a water box, while the proteins were minimized using ICM to optimize side chain orientations and hydrogen bonding with the water box. Then the water was deleted, as well as any salts from the crystal structure. The AcpP protein structures were obtained from the MD experiments, and were docked with the remaining substrate. No minimizations or solvation were performed on the AcpP structures, in order to retain the structure from the MD simulations. Subsequently the proteins were docked using the ICM1 Fast Fourier Transform docking method. The docking procedure was refined using previously identified key residues (Nguyen et al, 2014). For the FabA simulations we identified from crosslinking studies (Nguyen et al, 2014) a focus residues D38, E41, and E47 from AcpP and residues R132 and R136 from FabA. We specified focus residues in order to narrow the resultant structures, given the breadth of experimental data. For the FabF simulations the AcpP residues D35, and T39 were identified to be important to the interaction, while residues K65 of chain A and R206 of chain B were chosen for FabF (Mindrebo et al., 2020). During the docking simulations the partner proteins used were identical with the only difference being the conformation of the AcpP generated by the MD simulations. The docking results were ranked in order of total energy. The scores are presented as calculated by the ICM program, with the total energy term, electrostatic contribution, van der Waals energy, and surface term. The number of interacting residues was determined by collecting all atoms on the partner protein which are within 3.5 Å of the AcpP. All protein structure figures were generated in PyMOL (The PyMOL Molecular Graphics System, Version 2.0 Schrödinger, LLC)

## Results and Discussion

### Comparative Genomics of the parental (WT) and evolved strain (AN62)

In total 18 mutations were identified in the newly sequenced genomes of the WT parental and the AN62 evolved strains when compared to the reference genome for *E. coli* K-12 MG1655 (Table 3 and 4). The observed mutations included single nucleotide polymorphisms (SNP), insertions, and deletions. Out of the 18 mutations, 3 mutations were present in both the parental and evolved strain (Table 3), 3 mutations were only present in the parental strain, and 12 mutations were observed only in the evolved AN62 strain (Table 4).

**Table 3.** Mutations in both the parental (MG1655) and evolved (AN62) strain of *E. coli* K12.

| Gene | Function | Mutation | Gene Position<br>(in nucleotides) | Region | Genome<br>Position |
| --- | --- | --- | --- | --- | --- |
| *gatC* | galactitol-specific PTS enzyme | Deletion<br>ACC→A | 918 | Coding | 2173360 |
| *glpR* | DNA-binding transcriptional repressor | Insertion<br>C→CG | 150 | Coding | 3560455 |
| n/a | Intergenic region | Insertion<br>A→ACG | n/a | Repeat<br>region | 4296380 |

n/a, not applied

**Table 4.**
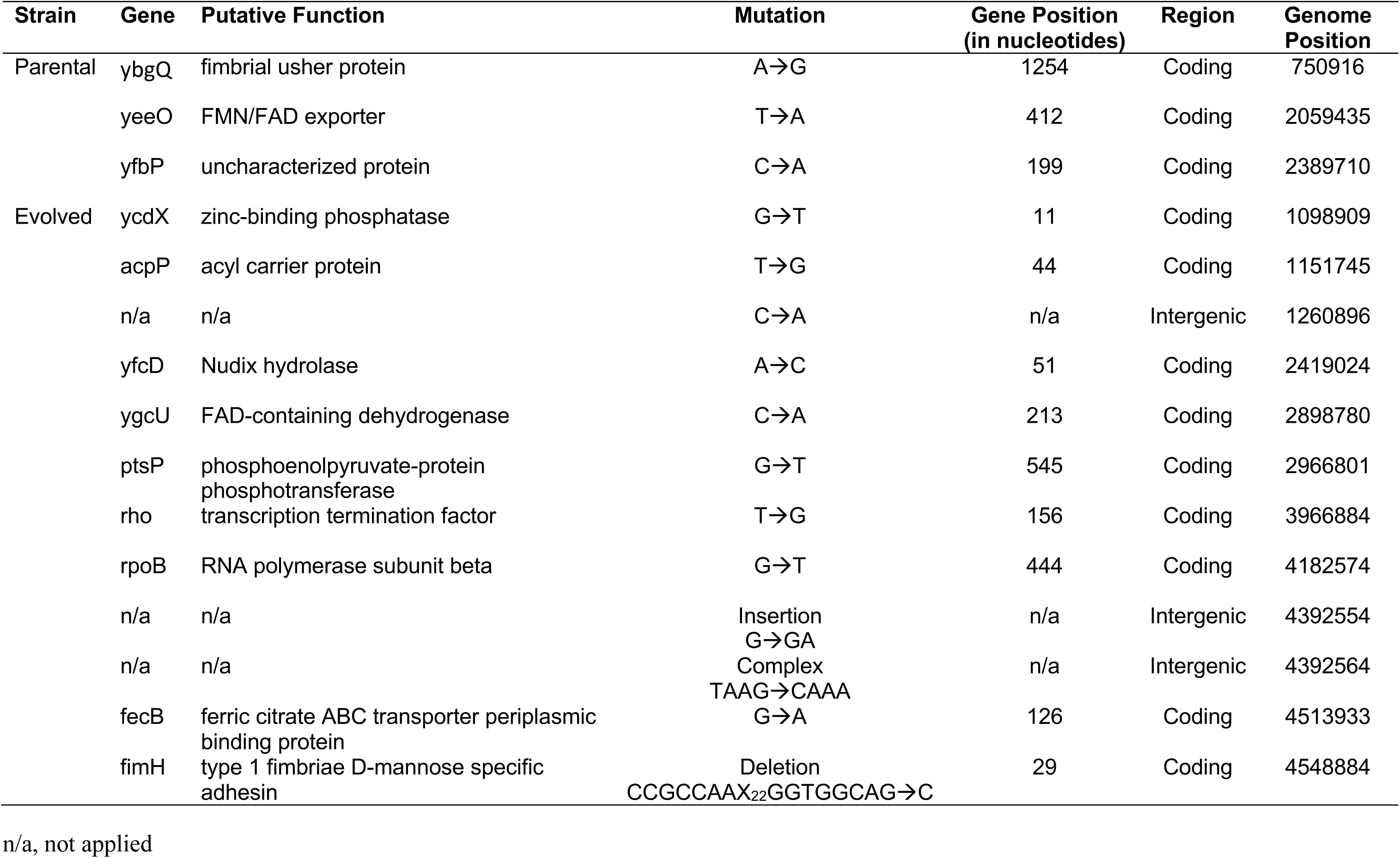
Strain specific mutations in the parental (MG1655) and evolved (AN62) strain of *E. coli* K12.

The mutations observed in both strains in comparison to the reference genome most likely arose before the start of the adaptive-laboratory-evolution experiment and was carried though the 500 generations of selection to the evolved strain. As demonstrated previously (Marietou et al., 2015), the parental strain did not exhibit altered growth phenotype at high pressure (60 MPa) and we are confident that the observed mutations are not linked with the ability of the evolved strain to grow at high pressure (60 MPa). The first mutation is caused by a deletion of two nucleotides in the coding sequence of *gat*C, a galacticol permease, resulting to a frameshift mutation (Table 3). The second mutation is caused by an insertion in the coding sequence of *glp*R, a transcription repressor, resulting again in a frameshift mutation (Table 3). The third mutation is an insertion of two nucleotides in a repeat region between *glt*P and *yj*cO genes (Table 3). The *gat*C and *glp*R mutations have been previously described in stock cultures of the *E. coli* K12 MG1655 strain (Freddolino et al., 2012). Domestication-related mutations such as the above are not uncommon in rapidly evolving bacterial strains such as *E.coli* (Freddolino et al., 2012).

Interestingly, we identified three additional parental strain specific mutations that were not present in the evolved strain (Table 4). The first mutation is caused by a single nucleotide substitution (SNP) in the coding sequence of *ybg*Q, a fimbrial usher protein, resulting in a synonymous mutation that does not affect the encoded amino acid (Table 4). The second mutation is a missence mutation affecting the structure of the flavin and dipeptide exporter encoding gene, *yee*O (Table 4). The third mutation is also a missence mutation in the coding sequence of an uncharacterized gene, *yfb*P (Table 4). Since these mutations are absent from the evolved strain we speculate that they were most likely reverted during the course of the ALE experiment. The reversion of the mutations in the pressure adapted AN62 strain could suggest that either these reversions resulted to an increased fitness under high pressure or they were associated with stochastic events.

### Evolved strain (AN62) associated mutations

Among the 12 unique mutations observed in the genome of the evolved strain, was a transversion mutation (T to G) in the *acp*P gene which encodes for the acyl carrier protein, as was previously described in our original ALE paper (Table 4) (Marietou et al., 2015). We speculated that the valine to glycine substitution (*acp*P^V43G^) increased the efficiency of unsaturated fatty acid synthesis contributing to the piezotolerant phenotype of the evolved strain. Given the essential nature of *acp*P and its importance in cell physiology this mutation was selected for further investigation (see relevant sections below).

We observed a missence mutation in the coding sequence of *ycd*X, resulting in a valine to phenylalanine (Val11Phe) amino acid change (Figure 1). YcdX belongs to the polymerase histidinol phosphatase protein superfamily and is involved in swarming motility as *ycd*X deletion mutants are unable to swarm (Redelberger et al., 2011). Motility is one of the most pressure-sensitive cell processes, affected by pressures as low as 10 MPa in *E.coli* (Bartlett 2002), thus the observed mutation could potentially be a direct result of the ALE at high pressure. Position V11 is variably conserved among well-studied piezophiles (Figure 1). Piezophilic *Psychromonas hadalis* K41G possess V11, while piezophilic *Photobacterium profundum* SS9 and *Shewanella benthica* KT99 have isoleucine at position V11 (Figure 1). The presence of isoleucine at position V11 for the piezosensitive *Photobacterium profundum* 3TCK suggests that modification of this residue alone does not equate to pressure adaptation.

**Figure 1.**
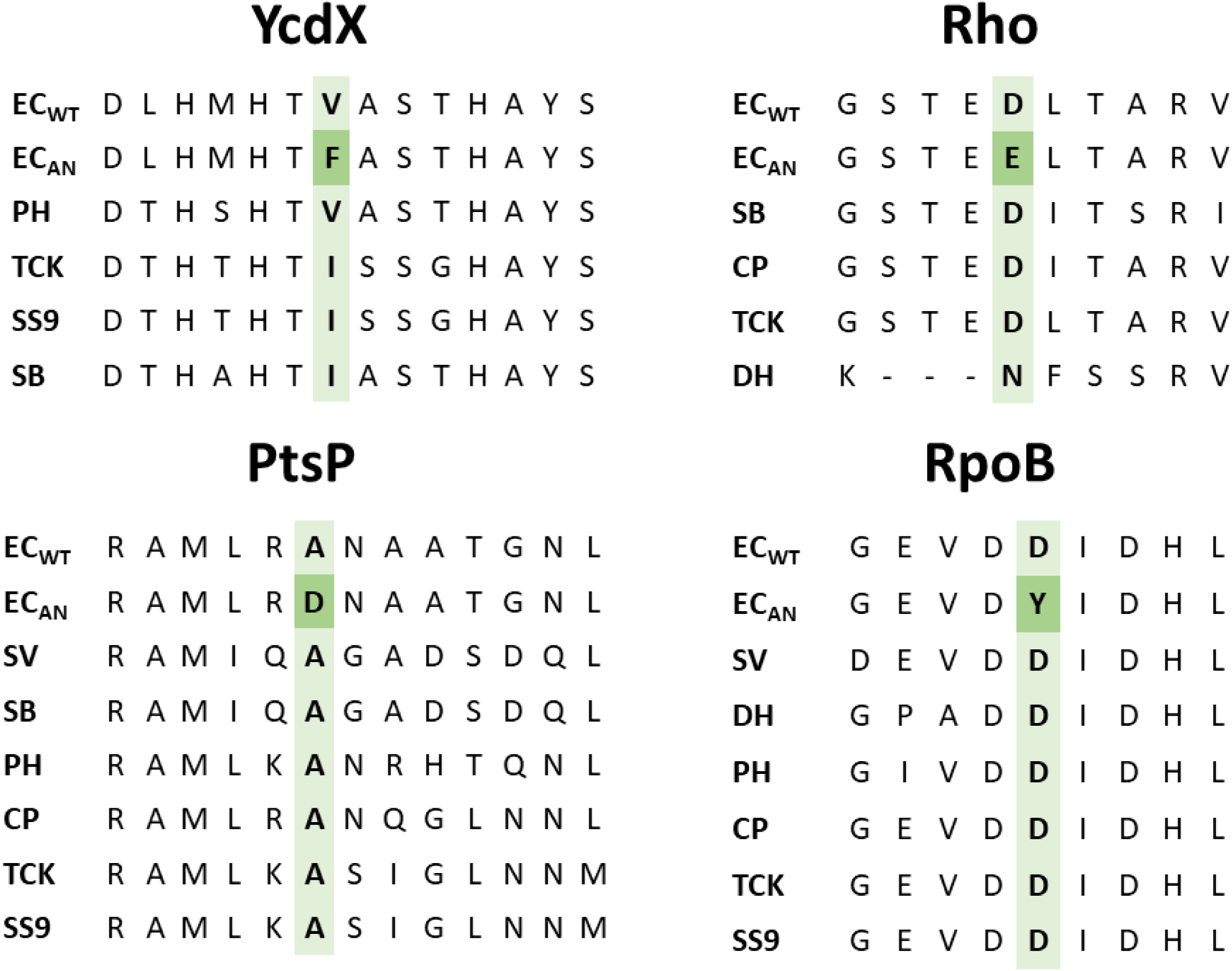
Alignment of protein sequences from the parental and evolved *E.coli* strains and other piezosensitive/piezophilic isolates. Highlighted in green are the residues of interest. ECWT, *E. coli* parental MG1655 strain; ECAN, *E. coli* evolved AN62 strain; PH, *Psychromonas hadalis*; TCK, *Photobacterium profundum* 3TCK; SS9, *Photobacterium profundum* SS9; SB, *Shewanella benthica* KT99; CP, *Colwellia piezophila*; DH, *Desulfovibrio hydrothermalis* AM13; SV, *Shewanella violacea* DSS12 .

We also observed SNPs in the coding sequence of a putative nudix hydrolase (*yfc*D), an uncharacterized FAD-linked dehydrogenase (*ygc*U), a phosphotransferase (*pts*P), a transcription termination factor (*rho*), the RNA polymerase beta subunit (*rpo*B), and a ferric citrate transporter (*fec*B) (Table 4). Transporters are greatly affected by high pressure since pressure affects reactions accompanied by large volume changes and was further confirmed in transcriptional studies (Simonato et al., 2006). We observed a missense mutation resulting in a leucine to phenylalanine (L126F) substitution in *fec*B, encoding for the periplasmic substrate-binding component of an iron citrate ABC transporter (Table 4). There was a positive selection for pressure based on amino acid substitution rates between deep-sea/piezophilic and shallow water/piezosensitive strains (Campanaro et al., 2008). We speculate that the observed substitution of leucine with the aromatic phenylalanine observed in the pressure-adapted *E. coli* could contribute to conformation stability of the transporter at high pressure.

The SNP within the coding sequence of *rho* resulted in the conservative substitution of aspartic to glutamic acid (D156E) (Figure 1). Even though conservative substitutions usually have less severe effects on the function of the protein, previous work in thermophiles suggested that substitution of aspartic acid with glutamic acid increased the transition temperature of the protein conferring higher stability (Lee et al., 2004). It is unclear whether such conservative amino acid substitution could also increase protein stability at high pressure conditions.

The SNP within the coding sequence of the FAD-linked dehydrogenase YgcU resulted in a missense mutation (Table 4). Previously, succinate dehydrogenase activity was significantly impacted in *E.coli* by pressures as low as 20 MPa while formic, and malic dehydrogenase activity was also affected by moderate pressures (Morita and Zobell, 1956; Morita, 1957). Saito et al. (2006.) reported several amino acid substitutions in the malate dehydrogenase of piezophilic isolates that might involved in pressure adaptation.

The *pts*P encodes for a phosphoenolpyruvate-dependent phosphotransferase with a SNP resulting in an alanine to aspartic acid (A545D) amino acid change in the evolved strain (Table 4, Figure 1). The A545 position is highly conserved among the examined piezophilic and piezosensitive genomes; however, that does not preclude that the observed mutation does not have an effect on the stability of the protein under increased pressure. PtsP activity requires the binding of a Mg^2+^ cofactor at the C-terminus (Rabus et al., 1999). Aspartic acid has a negatively charged side chain that assists in the binding of divalent cations such as Mg^2+^, thus we speculate that the observed substitution could be involved in stabilizing the cofactor binding under high pressure.

The SNP observed at the RNA polymerase b subunit-encoding gene, *rpo*B, resulted in a non-conserved aspartic acid to tyrosine (D444Y) amino acid substitution (Table 4, Figure 1). The effects of high pressure on the transcriptional activity of the RNA polymerase are noticeable from 50 MPa in *E.coli*, while in the piezophile *Shewanella violacea* the RNA polymerase was more active and stable at increasing pressure (Kawano et al., 2004). RNA polymerase is essential for transcription thus any SNP would will most likely have multiple effects on the physiology of the cell (Koch et al., 2014). Several studies have highlighted that *rpo*B mutations in *E. coli* are associated with altered physiology and adaptation to environmental stress (Herring et al., 2006; Rodríguez-Verdugo et al., 2013; Dragosits et al., 2013). An aspartic acid to tyrosine (D654Y) change within RpoB increased osmotolerance and succinic acid production in *E.coli* (Xiao et al., 2017). The D554Y most likely caused a conformation change that affected the transcriptional activity of RpoB and the upregulation of several osmotic response genes (Xiao et al., 2017). Similarly, we speculate that the D444Y mutation in RpoB could facilitate the pressure-adapted response (gene level) observed phenotypically in the pressure-adapted *E.coli* strain.

A disruptive in-frame deletion of 37 nucleotides from *fim*H, a type I fimbrin D-mannose specific adhesion encoding gene, was observed in the pressure adapted-strain that most likely resulted in loss of function (Table 4). Deletion of the *fim*H gene in E. coli did not affect the expression and morphology of type I pili since the encoded FimH adhesin forms only the tip of the pili affecting the length of the pili and the ability of E. coli to infect cells (Jones et al., 1995; Busch et al., 2015). Interestingly, the evolved strain lacked a parental strain SNP in the *ybg*Q gene encoding a fimbrial usher protein suggestive of a possible pressure adapted reversion mutation during the ALE experiment. Finally, we also identified an insertion and a complex nucleotide polymorphism in a non-coding region between two tRNA encoding genes (Table 4). The possible contribution of these mutations to the phenotype of the pressure-adapted strain is unclear.

### The effect of the *acp*P^V43G^ mutation on high-pressure growth and fatty acid production

A *ycf*H::cat mutation was introduced in the parental (MG1655) and the evolved AN62 strains using lambda red recombineering to generate the strains MAE34 (MG1655 *ycf*H::cat) and MAE35 (AN62 *ycf*H::cat). Once verified, P1 lysates of these strains were used to cross the linked *acp*P allele into either MG1655 or AN62 strains. The resulting strains MAE36 (MG1655 *acp*P^V43G^) and MAE37 (AN62 *acp*P^WT^) were further examined for their high-pressure growth phenotype and their membrane fatty acid composition. The growth phenotype of the engineered strains (MAE36 and MAE37) was compared to those of the parental (MG1655) and evolved (AN62) strains (Figure 2). The pressure evolved strain AN62 grew well at 60MPa as indicated by the color change of the culture medium to violet due to microbial growth, while no growth was observed for MG1655 following 48hr of incubation (Figure 2). Similar to the parental strain, MAE36, an MG1655 derivative containing the *acp*P^V43G^ mutation, displayed no growth at 60 MPa (Figure 2). Strain MAE37, an AN62 derivative with the *acp*P^WT^ allele grew at high pressure similarly to the evolved AN62 strain (Figure 2). Three independently derived transductants were assessed for each strain with no variation noted in the growth phenotype (Figure 2). Our results suggest that the V43G mutation found in *acp*P is not solely responsible for the respective high-pressure growth phenotypes found in the pressure-adapted AN62 strain.

**Figure 2.**
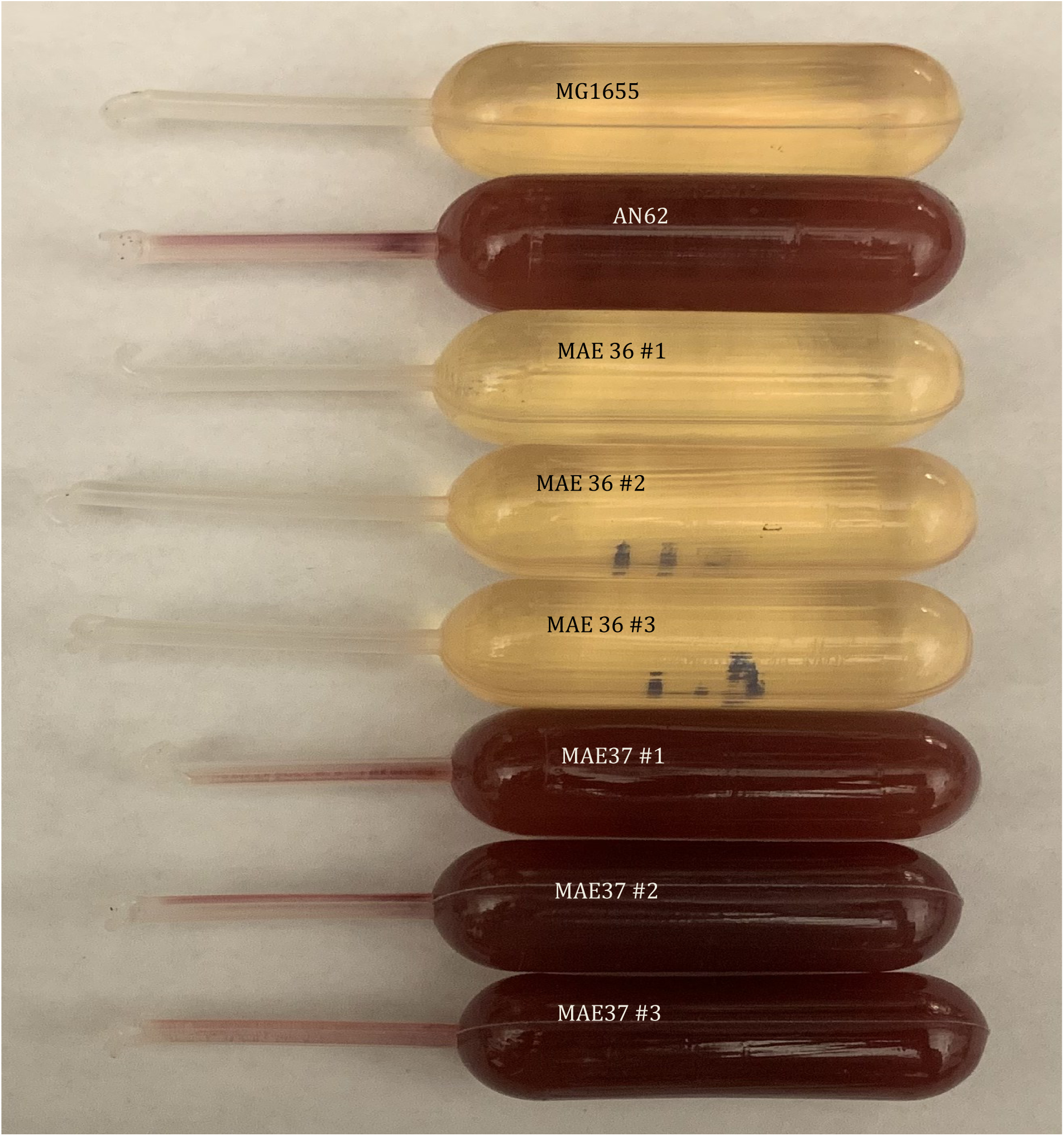
High pressure (60 MPa) growth of the parental, pressure adapted, and engineered *E. coli* strains after 48 h of incubation at 37°C. Tetrazolium chloride (TTC) was added as colorimetric growth indicator. MG1655, parental strain; AN62, pressure adapted strain; MAE36, MG1655 acpP^V43G^ strain; MAE37, AN62 acpP^WT^.

As described previously (Marietou et al., 2015), we confirmed once again that the pressure adapted AN62 strain contained a significantly higher amount of cis-vaccenic acid (18:1) content relative to the parental MG1655 strain while the overall unsaturated to saturated fatty acid ratio (UFA/SFA) remained the same (Table 5). Compared to the parental strain, the fatty acid profile of MAE36 showed a noticeable increase in the abundance of cis-vaccenic acid (18:1) with minor variations in the other fatty acid species (Table 5). A comparison of AN62 and MAE37 fatty acid profiles showed a decrease in cis-vaccenic acid content in MAE37, which led to an overall decrease in the UFA/SFA ratio (Table 5). These results provide further evidence that the V43G mutation is associated with an altered fatty acid profile and in particular with increased levels of cis-vaccenic acid. Earlier work in E.coli has demonstrated that FabF is able to regulates membrane fluidity by converting cis-palmitoleoyl-AcpP (16:1-AcpP) into cis-vaccenoyl-AcpP (18:1-AcpP) in response to temperature changes (Garwin and Cronan, 1980). Given that alterations of cis-vaccenic acid (18:1) are associated with the presence or absence of the V43G mutation, this mutation is likely involved with modulation of FabF activity. To the best of our knowledge, this work demonstrates the first example of a discrete mutation in *acp*P being responsible for an altered fatty acid profile in *E. coli*.

**Table 5.** Fatty acid profiles of the of the parental (MG1655), pressure adapted (AN62), and engineered (MAE36/MAE37) *E.coli* strains grown aerobically at 37°C.

|  | Mean % (by weight) fatty acid<sup>a</sup> |  |  |  |
| --- | --- | --- | --- | --- |
| Fatty acid | MG1655 | AN62 | MAE36 | MAE37 |
| 12:0 | 3.75 ± 0.27 | 3.87 ± 0.05 | 3.59 ± 0.21 | 3.91 ± 0.14 |
| 14:0 | 9.66 ± 0.23 | 5.90 ± 0.25 | 6.50 ± 0.32 | 7.88 ± 0.16 |
| 14:0 3OH | 4.72 ± 0.66 | 5.07 ± 0.75 | 6.36 ± 1.48 | 6.10 ± 0.16 |
| 16:0 | 40.63 ± 2.07 | 41.56 ± 1.54 | 32.71 ± 0.90 | 46.42 ± 0.79 |
| 16:1 | 25.43 ± 0.40 | 14.22 ± 0.64 | 26.95 ± 3.33 | 15.28 ± 0.02 |
| 18:1 | 14.72 ± 0.40 | 24.60 ± 0.66 | 22.88 ± 0.43 | 16.59 ± 0.07 |
| 18:0 | 0.32 ± 0.45 | 0.76 ± 0.65 | 0.00 ± 0.00 | 0.73 ± 0.63 |
| UFA/SFA | 0.68 | 0.68 | 1.01 | 0.49 |

<sup>a</sup> Data represents the average of at least three independent replicates; ± standard deviation

### Investigating the effect of the *acp*P^V43G^ mutation using molecular dynamics (MD) simulations

To investigate the structural rationale of the V43G AcpP phenotype, we performed MD simulations of the WT and V43G AcpP protein harboring two different acyl substrates. Three independent simulations were carried out each for 500 ns. Two critical steps are important for unsaturation in the fatty acid biosynthesis pathway. First, dehydration of 3R-hydroxydecanoyl-AcpP to 2E-decanoyl-AcpP and subsequent isomerization of 2E-decanoyl-AcpP to 2Z-decanoyl-AcpP by FabA commits the unsaturation into the FAS pathway by preventing further reduction. (Heath and Rock 1996) Second, elongation of 9Z-hexadecenoyl-AcpP to 11Z-octadecenoyl-AcpP by FabF commits 11Z-octadecenoic acids to the *E. coli* membrane. Therefore, the AcpP was simulated with either 3R-hydroxydecanoyl- or 9Z-hexadecenoyl-groups bound. Throughout the simulations, the acyl chain and phosphopantetheine group sampled dynamic states, transitioning between elongated and kinked conformations. This kinked conformation has been observed in MD simulations carried out by Chan et al. (2008) and described as a sub-pocket within AcpP mediated by a conformation switch of χ1 angle of residues L42 and L46. Surprisingly, the V43G mutation resulted in increased stability of the elongated 3R-hydroxydecanoyl-chain compared to the WT 3R-hydroxydecanoyl-AcpP and V43G and WT 9Z-hexadecenoyl-AcpP (SI figures). The V43G mutant did not have a large effect on the overall backbone dynamics in the 3R-hydroxydecanoyl-AcpP, however the V43G mutant 9Z-hexadecenoyl-AcpP was much less dynamic than the WT in all regions except for loop II and helix III of the conserved AcpP binding site (Figure 3). A larger mean pocket volume was calculated for the V43G AcpP simulations compared to the WT AcpP, which was more pronounced in the 9Z-hexadecenoyl-AcpP simulation (Figure 4).

**Figure 3.**
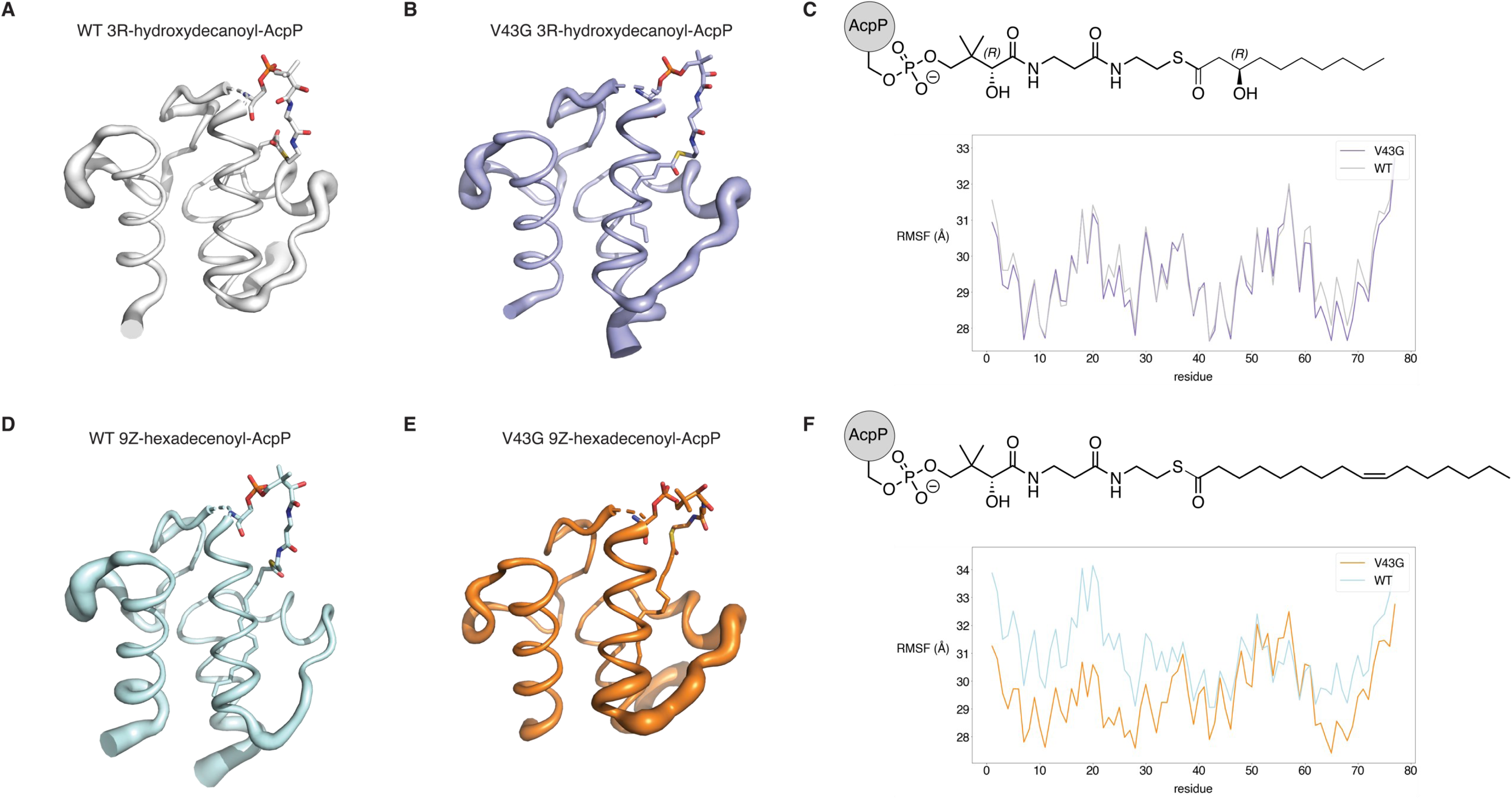
Root mean squared fluctuation (RMSF) during molecular dynamics simulations of WT and V43G AcpP. (A-C) 3R-hydroxydecanoyl-AcpP with cartoon thickness based on RMSF per residue calculated from 1.5 μs aggregate simulation of WT (A), and V43G (B), shown graphically underneath structure schematic in (C). (D-F) 9Z-hexadecenoyl-AcpP with cartoon thickness based on RMSF per residue calculated from 1.5 μs aggregate simulation of WT (D), and V43G (E), shown graphically underneath structure schematic in (F).

**Figure 4.**
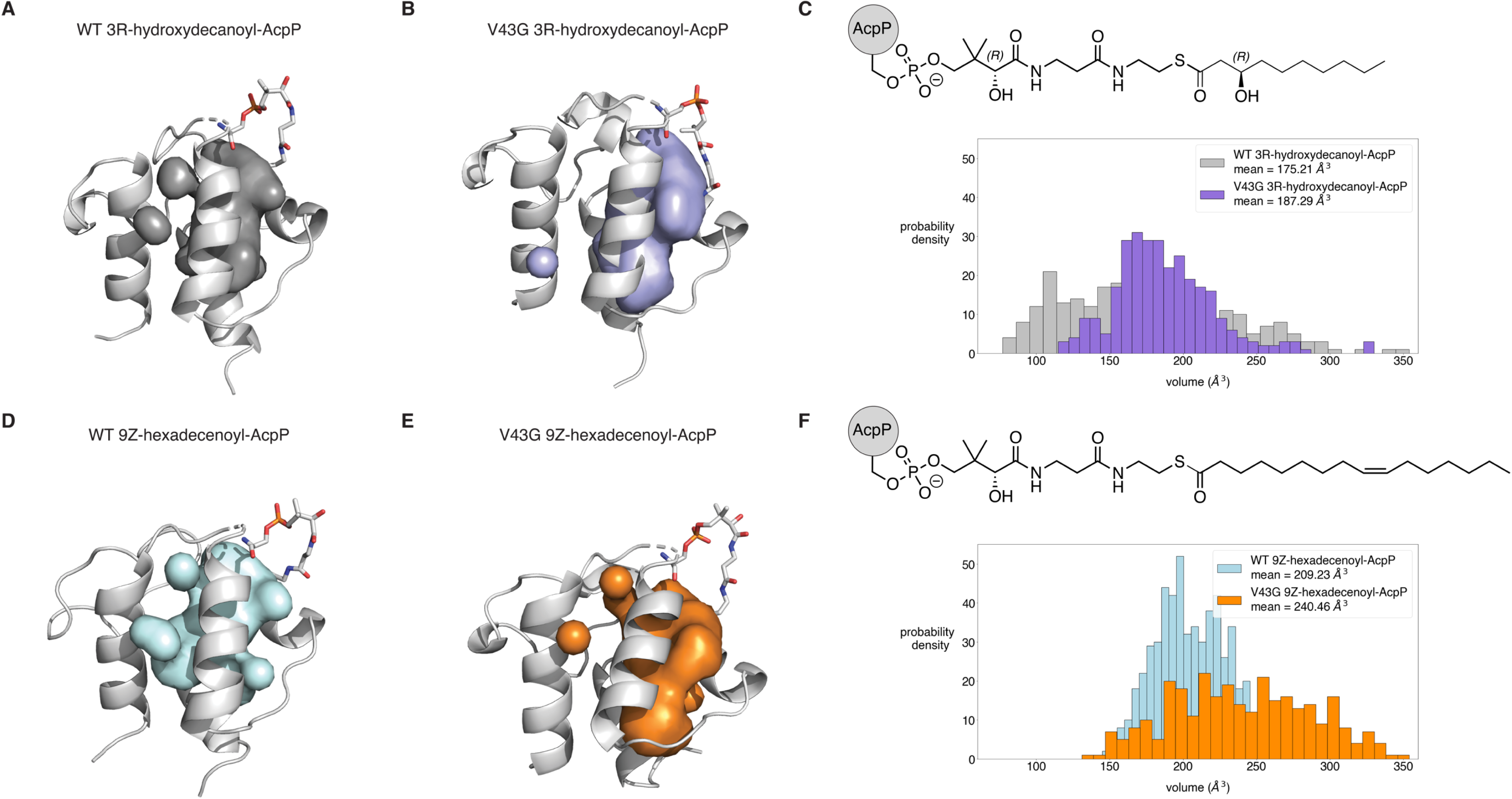
Pocket volume calculated from molecular dynamics simulations of WT and V43G AcpP. (A-C) 3R-hydroxydecanoyl-AcpP with pocket volume shown for WT (A), and V43G (B). Histogram showing the distribution of pocket volume calculated from 1.5 μs aggregate simulation underneath structure schematic in (C). (D-F) 9Z-hexadecenoyl-AcpP with pocket volume shown for WT (D), and V43G (E). Histogram showing the distribution of pocket volume calculated from 1.5 μs aggregate simulation underneath structure schematic in (F).

Interestingly, mutations of residue V43 in AcpP of *E.coli* have been previously noted for a variety of different phenotypes (Angelini et al., 2012; Keating and Cronan, 1996). A V43I mutation was previously shown to be associated with increased electrophoretic mobility in native acrylamide gels suggesting a compact structure (Keating and Cronan, 1996). A two-hybrid screen for mutations that impact the ability of AcpP to interact with SpoT revealed a V43E mutation that impaired this interaction (Angelini et al., 2012). Structurally, the V43 residue is found in helix II of AcpP, which has been shown to be the interface site of multiple AcpP interactions (Finzel et al., 2015; Zhang et al., 2001) that could potentially affect the interaction of AcpP with enzymes from the FAS pathway.

### Docking simulations of the V43G AcpP protein to FabA and FabF

In order to test whether the structural differences identified in the MD analysis translate to increased binding affinity for the enzymes responsible for unsaturation, we performed docking simulations of the WT and V43G AcpP proteins to FabA and FabF. Unsaturations are first installed when C10:1 AcpP is isomerized by the FabA dehydratase enzyme. In order to test if the V43G mutation increases the likelihood of this event, high resolution docking studies were performed to evaluate the binding of WT and mutant AcpP. AcpPs from the most populated cluster of the WT and V43G MD simulation were docked to FabA, the outputted conformations were sorted by the most stable and the top three structures were examined further. All three structures were able to dock within 5Å of the crosslinked interface; however the V43G AcpP structures exhibited stronger binding compared to the WT counterparts (Figure 5). The relative energetics as demonstrated by the electrostatic contributions to binding (Van der Waal’s forces, surface area, and overall energetics) suggest that the mutant V43G AcpP structures form a stronger interface (Figure 5).

**Figure 5.**
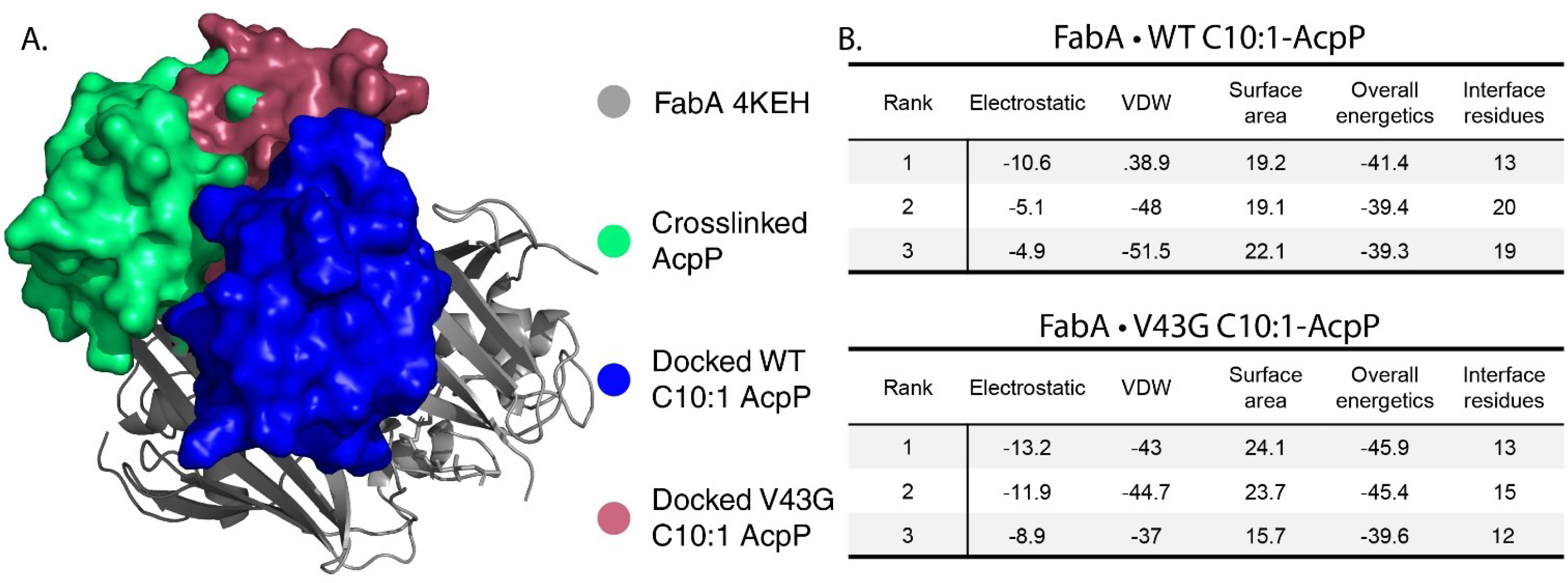
A) The structure of FabA crosslinked AcpP (pdb:4KEH) in comparison with docked WT and V43G C10:1 AcpP. The 2 docked structures bind slightly different surfaces of FabA. B) The docking energetics, surface area, and interacting residues of the top 3 structures of WT and V43G C10:1 AcpP with FabA. These data have demonstrated that V43G AcpP binds with a stronger interface, especially with a much stronger electrostatic interface in comparison to WT C10:1 AcpP.

The second step in the creation of C18:1 fatty acids is the elongation from 16:1 to 18:1 by the FabF ketosynthase (Beld et al., 2015). In order to test if the V43G mutation increased the affinity for FabF a second round of docking was performed from the C16:1 most populated clusters of AcpP with the C16 crosslinked FabF. Once again, the top three conformations were investigated. The docking simulations demonstrated that the V43G AcpP mutant bound FabF with a stronger interface, however the means of their binding differed between the WT and the AcpP mutant (Figure 6). While the WT FabF complex tended to bind with a larger interface and more total residues, the V43G AcpP mutant appeared to have a stronger interface, with consistent size, electrostatic contribution, and Van der Waal’s contributions to binding (Figure 6). These simulations offer further evidence that the V43G AcpP mutant forms stronger interactions with the enzymes responsible for installing and elongating fatty acid unsaturations.

**Figure 6.**
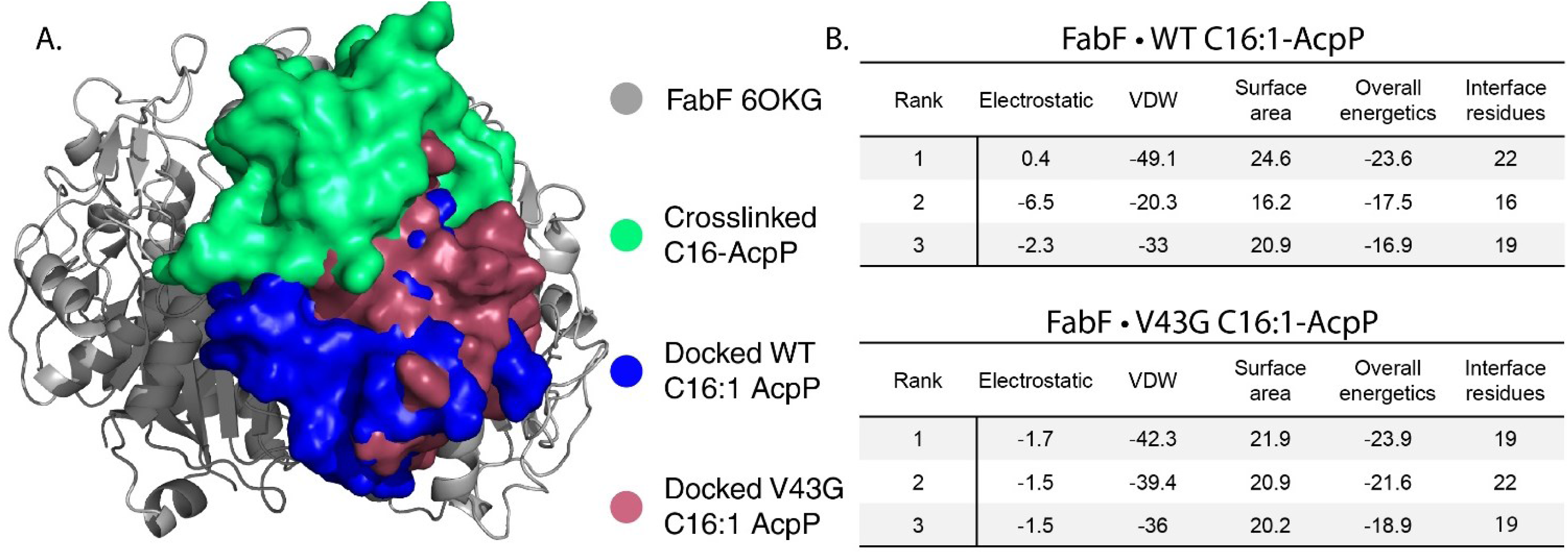
A) The structure of FabF crosslinked AcpP (pdb:6OKG) in comparison with docked WT and V43G C16:1 AcpP. The 2 docked structures overlay very closely with one another and the crosslinked structure. B) The docking energetics, surface area, and interacting residues of the top 3 structures of WT and V43G C16:1 AcpP with FabF. These data demonstrate that although the two docked structures bind at very similar interfaces the V43G mutant binds with stronger affinity.

## Conclusion

There is no definitive list of genes necessary for microbial life at high pressure; this study offers a glimpse of genome level adaptations that have the potential to contribute to increased pressure tolerance. Even though the V43G mutation in AcpP is not responsible for high-pressure growth adaptation in the evolved AN62 strain, it is associated with increased levels of cis-vaccenic acid and with an altered fatty acid profile. MD and docking simulations suggest that the V43G mutation affects the interaction of the AcpP protein with other enzymes of the fatty acid biosynthetic pathway such as FabA and FabF promoting a stronger interaction. Further molecular studies in *E.coli* where wild-type genes will be replaced with the mutated versions as they appear in the evolved strain genome, would allow us to confirm the contribution of each individual mutation to the pressure-adapted phenotype. The list of mutated genes in the pressure-adapted strain was relative low suggesting that pressure-adaptation was most likely due to the combined effect of the observed mutations and substantial metabolic reprogramming (gene regulation). It would thus be interesting to study the gene expression profile (transcriptomics) of the evolved strain at atmospheric and high pressure (60 MPa) in future experiments.

## Acknowledgements

The authors would like to thank Anne Stentebjerg, Britta Poulsen, and Lykke Beinta Bjærge Bamdali for technical assistance.

## Author Contributions

Conceptualization, MNA, DHB, AM; Experimental procedures, MNA, SM, AM; molecular dynamics simulations, TS, TGB; bioinformatic analysis, IPGM; manuscript preparation-original draft, MNA, TS, SM, TGB, AM; manuscript preparation-review and editing, MNA, TS, TGB, IPGM, DHB, AM; funding, TS, IPGM, EEA, MB, JAM, AM.

## Ethical statement

No human nor animal experimentation is reported

## Data bibliography

