## Supplementary figures for "Adaptation to high pressure; insights from the genome of an evolved *Escherichia coli* strain with increased piezotolerance"

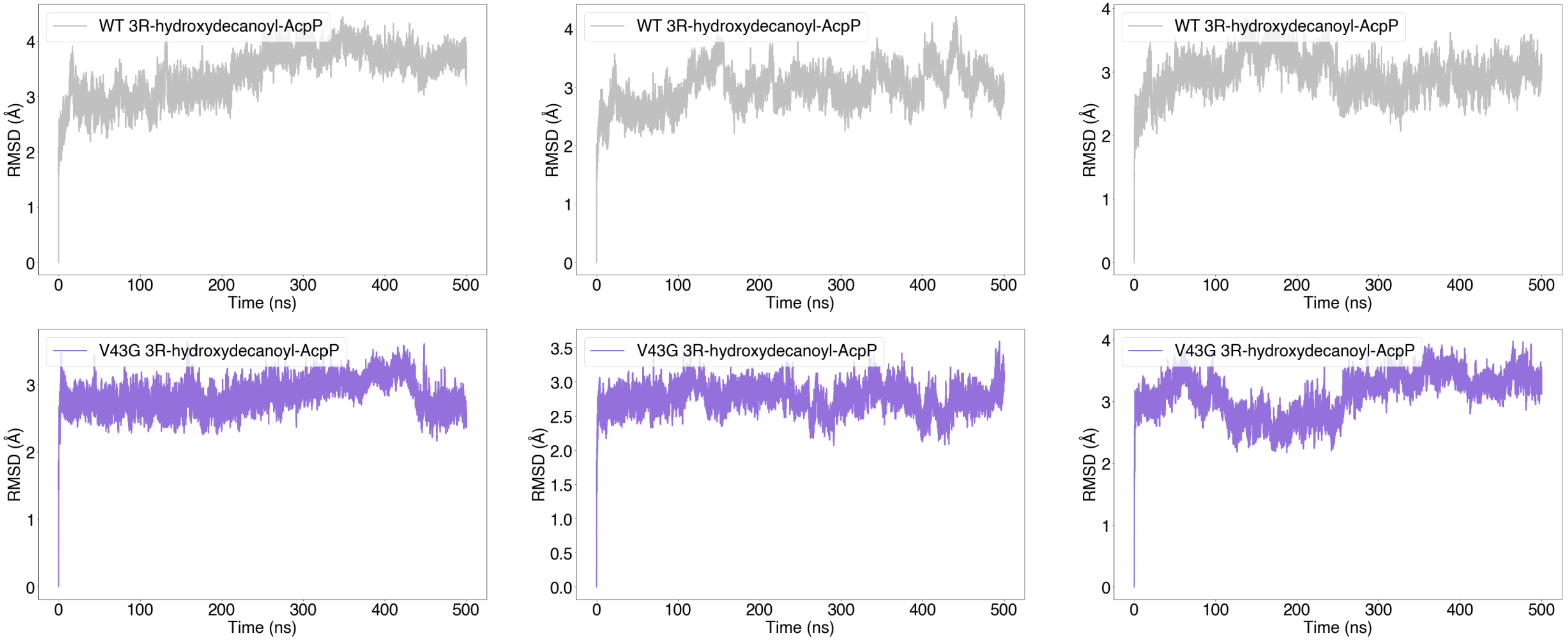


**Supplementary Figure 1.** All atom RMSD vs Time for each of three replicates simulated of WT 3R-hydroxydecanoyl-AcpP (top) and V43G 3R-hydroxydecanoyl-AcpP (bottom).

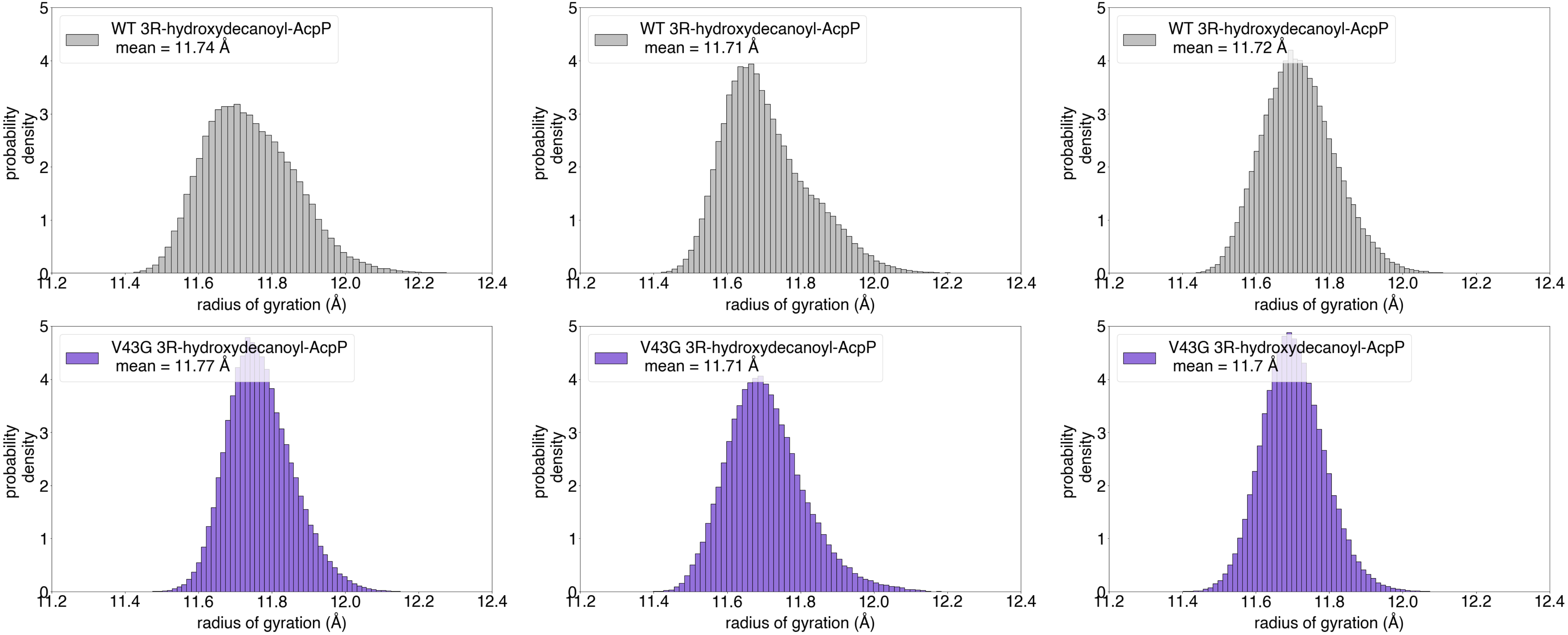


**Supplementary Figure 2.** Histograms representing the radius of gyration for each of three replicates simulated of WT 3R-hydroxydecanoyl-AcpP (top) and V43G 3R-hydroxydecanoyl-AcpP (bottom).

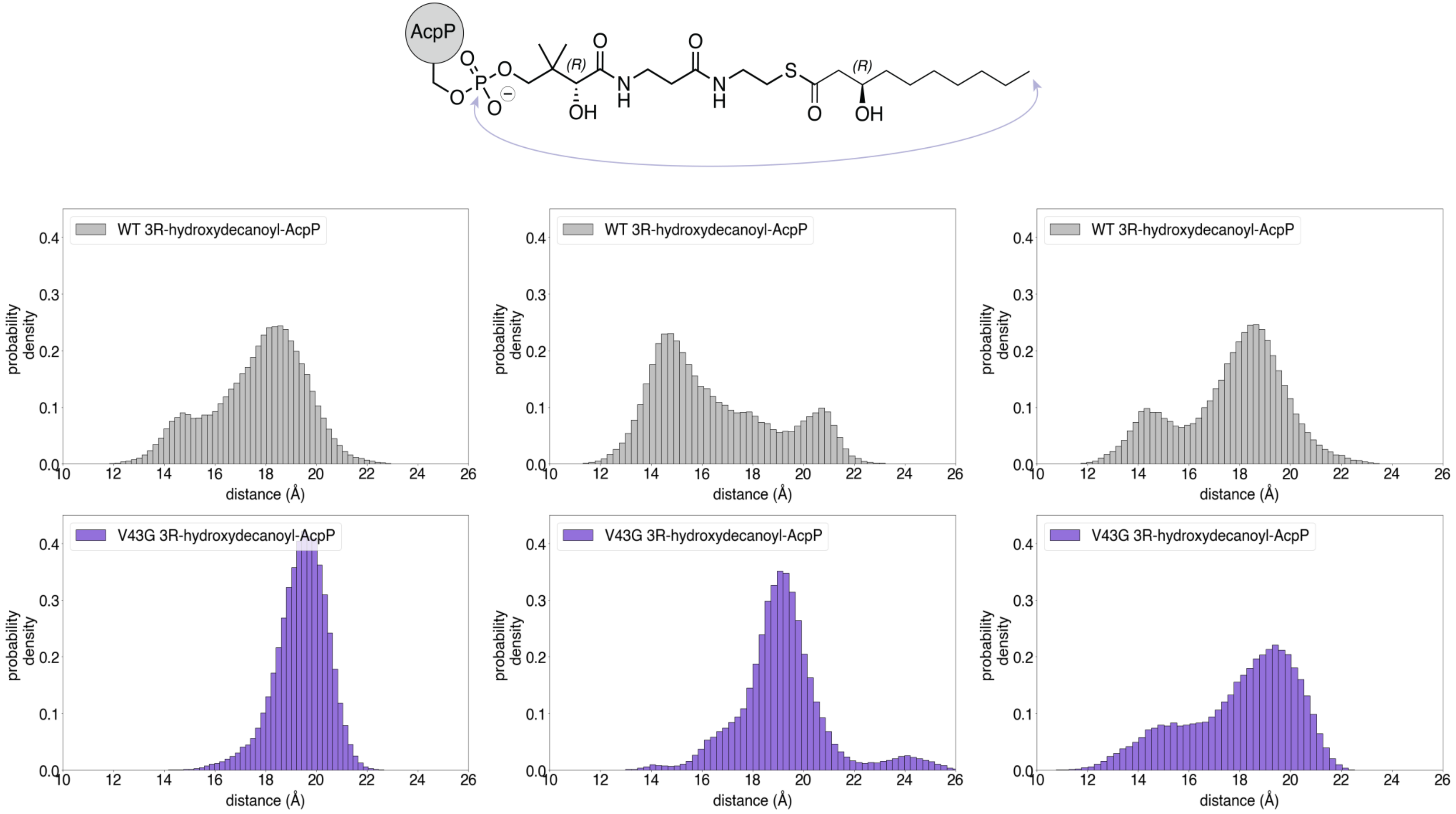


**Supplementary Figure 3.** Histograms representing the “elongated” versus “kinked” form of the ligand determined by calculated the distance between the phosphate and terminal carbon of the acyl chain for each of three replicates simulated as shown above for WT 3R-hydroxydecanoyl-AcpP (top) and V43G 3R-hydroxydecanoyl-AcpP (bottom).

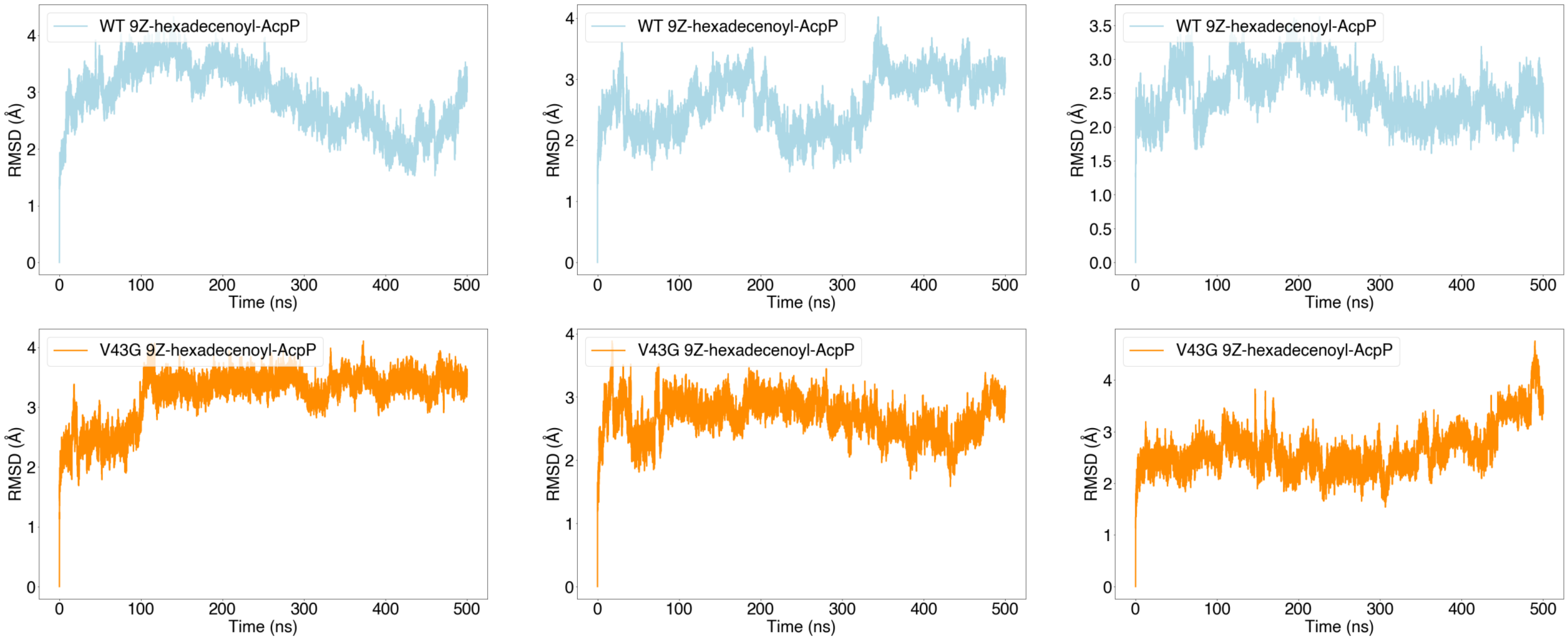


**Supplementary Figure 4.** All atom RMSD vs Time for each of three replicates simulated of WT 9Z-hexadecenoyl-AcpP (top) and V43G 9Z-hexadecenoyl-AcpP (bottom).

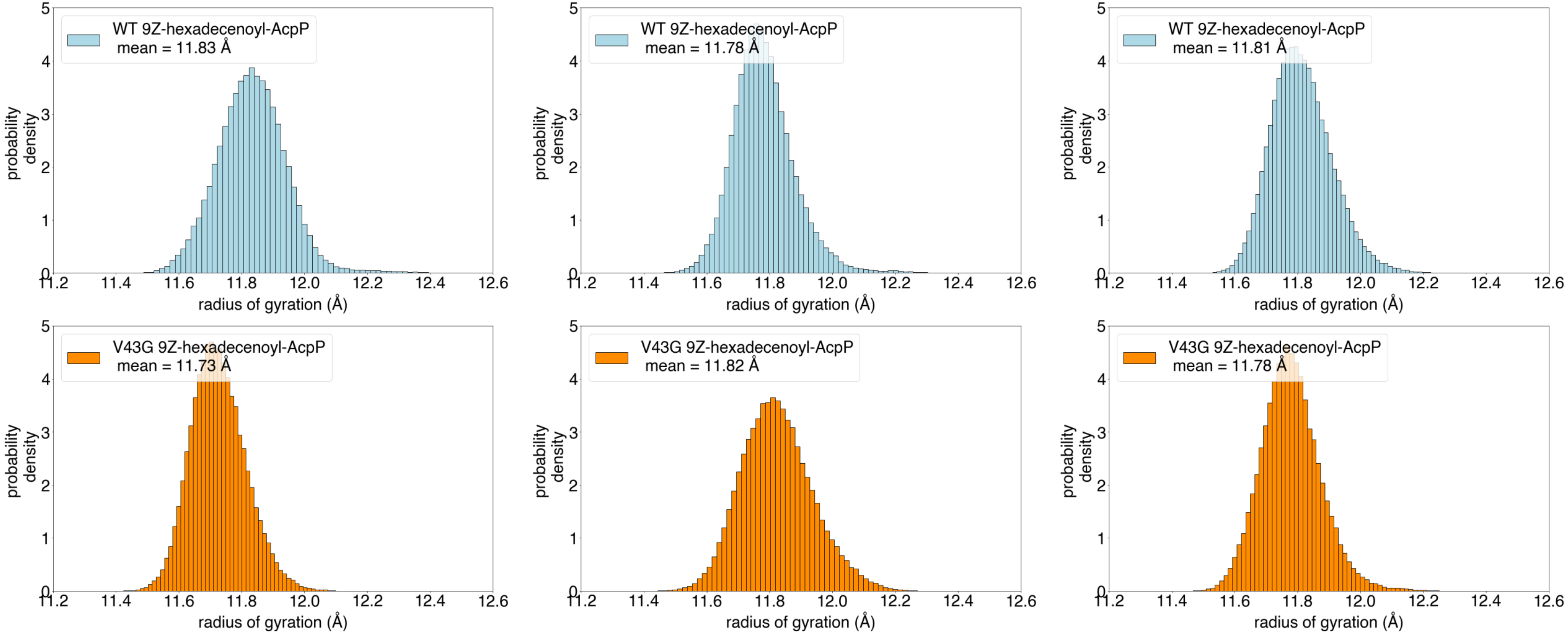


**Supplementary Figure 5.** Histograms representing the radius of gyration for each of three replicates simulated of WT 9Z-hexadecenoyl-AcpP (top) and V43G 9Z-hexadecenoyl-AcpP (bottom).

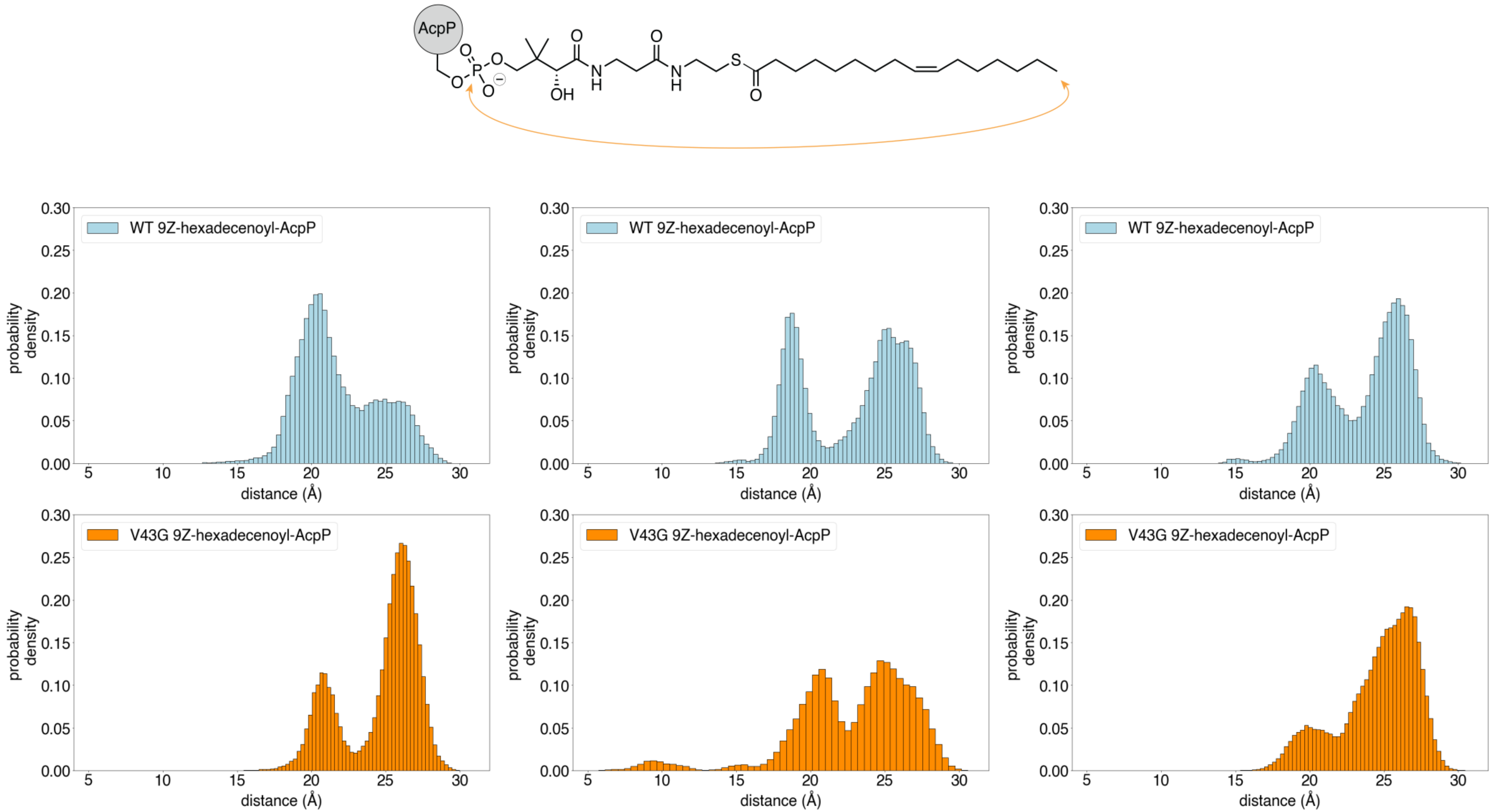


**Supplementary Figure 6.** Histograms representing the "elongated" versus "kinked" form of the ligand determined by calculated the distance between the phosphate and terminal carbon of the acyl chain for each of three replicates simulated as shown above for WT 9Z-hexadecenoyl-AcpP (top) and V43G 9Z-hexadecenoyl-AcpP (bottom).
